# Development and evaluation of a core genome multi-locus sequence typing scheme for the *Enterobacter cloacae* complex

**DOI:** 10.64898/2026.09.14.751580

**Authors:** Hilary Miller, Sarah Bakker, Kristin Dyet, David Winter

## Abstract

The *Enterobacter cloacae* species complex (ECC) comprises a group of closely related, opportunistic Gram-negative bacteria of major public health concern due to their frequent involvement in healthcare-associated infections and their capacity to acquire and disseminate multidrug resistance, including carbapenemases. Because members of this complex are often difficult to distinguish phenotypically and their taxonomic status is subject to debate, there is a need for a standardized, species-complex wide typing scheme based on whole genome sequencing (WGS) that can be applied to any species within the complex. In this study we have developed and evaluated a core genome multi-locus sequence typing (cgMLST) scheme suitable for species within the ECC. Using 3442 publicly available genomes from 27 ECC species or subspecies we developed a scheme with 1812 loci, comprising loci present in 99% of all genomes. Among the 3442 isolates in our study, 99.9% had 95% or more of the cgMLST targets, indicating that the schema is well-defined and representative for the breadth of ECC species in our study. On two independent evaluation datasets, the scheme reliably resolved epidemiologically linked isolates with 0–3 allelic differences and returned the same outbreak clusters defined previously by higher-resolution core-genome SNP (cgSNP) analysis. Hierarchical clustering analysis at different levels of resolution showed that the cgMLST profiles could potentially be used to differentiate between species and sub-lineages in the complex. The cgMLST schema will improve the ability of public health laboratories to perform WGS-based surveillance of ECC species.

**Data Summary:** NCBI accession numbers for genome assemblies used to create the cgMLST schema are given in Supplementary Table 1. Samples used to evaluate the schema are from NCBI Bioprojects PRJEB46479 (1) and PRJNA1469251 (sequenced by PHF Science as part of this study). The cgMLST schema created in this study is publicly available from https://chewbbaca.online/species/21.

**Impact Statement:** The *Enterobacter cloacae* complex (ECC) is an increasingly important group of opportunistic, multidrug-resistant pathogens implicated in healthcare-associated infections worldwide, yet its surveillance is hindered by taxonomic complexity and the lack of standardized, scalable typing approaches. In this study, we present a robust, species-complex–wide core genome MLST (cgMLST) scheme derived from over 3,400 genomes spanning 27 ECC taxa, providing a unified framework that captures genomic diversity while maintaining high resolution for outbreak detection. The scheme performs consistently across species boundaries, overcomes limitations of SNP-based approaches for large and heterogeneous datasets, and demonstrates concordance with established outbreak definitions. By enabling portable, reproducible, and high-resolution genomic comparisons without requiring precise species identification, this resource directly addresses a critical gap in ECC genomic epidemiology. Its adoption will enhance the ability of public health laboratories to conduct real-time surveillance, detect transmission events, and respond effectively to the spread of antimicrobial-resistant ECC strains.

## Introduction

Species within the *Enterobacter cloacae* complex (ECC) are increasingly important nosocomial pathogens and are significant contributors to the global burden of healthcare-associated infections (2). The complex currently comprises more than 20 species or subspecies within the *Enterobacter* genus (3), a group of facultatively anaerobic, Gram-negative bacteria belonging to the family *Enterobacteriaceae*. Although members of the ECC are commonly found in the environment and form part of the commensal microbiota of the human and animal gut, they are also known to be opportunistic pathogens capable of causing a range of clinically significant infections including pneumonia, septicaemia and urinary tract infections (4).

ECC species are included within the ESKAPEE group of pathogens (*Enterococcus faecium, Staphylococcus aureus, Klebsiella pneumoniae, Acinetobacter baumannii, Pseudomonas aeruginosa, Enterobacter* spp., and *Escherichia coli*), which exhibit multidrug resistance and together represent the leading causes of nosocomial infections worldwide (5). ECC strains frequently harbour extended-spectrum β-lactamases (ESBLs) and carbapenemases, which are often co-located on plasmids with genes conferring resistance to other classes of antibiotics such as aminoglycosides and fluoroquinolones (4, 6-8), complicating therapeutic management.

Improved understanding of the genomic diversity and relatedness of ECC strains is essential for monitoring transmission events and informing infection prevention strategies. ECC species are difficult to distinguish using phenotypic methods commonly employed in clinical laboratories such as Gram staining, biochemical assays, and mass spectrometry (9). Despite the widespread use of whole genome sequencing (WGS) for distinguishing between ECC species (10-12), few tools exist for typing isolates for outbreak investigation. Core genome SNP-based methods (cgSNP) are often regarded as the gold standard for high resolution outbreak tracing and have been used for tracking transmission of ECC isolates in previous studies (1, 13). However, SNP based methods do not scale efficiently to large surveillance datasets, especially when samples may be from different members of a species complex. Core genome SNP pipelines typically require a closely related reference genome and use of a single reference across multiple species within a complex is likely to result in reduced core-genome size, and inaccurate SNP calling for more divergent strains (14).

Core genome multi-locus sequence typing (cgMLST) methods avoid many of the problems of SNP-based typing by using standardized, allele-based schemes that scale well to large, diverse datasets, and are reproducible and portable across different laboratories (15). The cgMLST schemes typically comprise a few thousand loci enabling greater discriminatory power than classical 7-gene MLST schemes, and are now a standard tool for surveillance of many pathogens of public health importance (16-19). CgMLST schemes are normally developed for individual species, but *ad hoc* schemes developed for individual ECC species in previous studies generally have not performed well in correctly identifying transmission events (20, 21), and are not suitable for applying more widely across the species complex. A publicly available scheme for *Enterobacter hormaechei* was recently developed but has not been tested for use against other species of ECC (22). There is a need for a standardized, species-complex wide schema for ECC as many public health surveillance programs rely on inaccurate lab methods for initial typing of ECC species and may incorrectly label isolates as *E. cloacae*, with the correct species uncertain until after WGS is performed. In addition, the taxonomic status of several species, including the most common clinical species *E. hormaechei* and its subspecies, is under dispute (10, 12, 23) complicating the development of single-species schema.

Development of a species-complex wide schema requires the inclusion of sufficient shared target loci to have enough resolution for tracking transmission events, while excluding loci that are not shared between the majority of species in the complex. Thus, species complexes with too much genomic diversity may lack enough shared loci to produce a useful schema. Species complex-wide schema have been successfully developed and applied to other species complexes such as *Klebsiella oxytoca/grimontii/michiganensis/pasteurii* (24) and *Citrobacter freundii/portucalensis/braakii/europaeus* (25) where the species are closely related and difficult to distinguish phenotypically.

In this study we aimed to develop a cgMLST schema that can be applied to all species within the ECC. To ensure accurate core genome representation across all members of the species complex we used a large number of input genomes from globally distributed sources, encompassing 27 different species or subspecies within the complex, to construct the schema. We assessed the suitability of the schema for identifying outbreaks in multiple different species of ECC and determined an allele distance threshold suitable for inferring transmission events.

## Methods

### ECC species

Sequences from 27 different species or subspecies of *Enterobacter* were included in our study (Table 1). In this study we have referred to all taxa by the valid species or subspecies names on the List of Prokaryotic names with Standing in Nomenclature (LPSN, (3)), which lists *E. hormaechei* with five distinct subspecies and *E. cloacae* with two subspecies. However, we note that there has been considerable recent debate around the taxonomy of the *E. hormaechei* and *E. cloacae* subspecies. Five subspecies of *E. hormaechei* are currently recognised (*E. hormaechei* subspecies *hoffmannii, hormaechei, oharae, steigwaltii* and *xiangfangensis*) (23, 26), but recent work based on WGS data has suggested that *E. hormaechei* subsp. *hoffmannii* should be elevated to a distinct species *E. hoffmannii*, and the *E. hormachei subspecies oharae, steigwaltii* and *xiangfangensis* should be subspecies under a new species *Eneterobacter xiangfangensis* (6, 10). It has also been proposed that the two subspecies of *E. cloacae* subsp. *cloacae* and *E. cloacae* subsp. *dissolvens*, should be separate species (10).

**Table 1.**
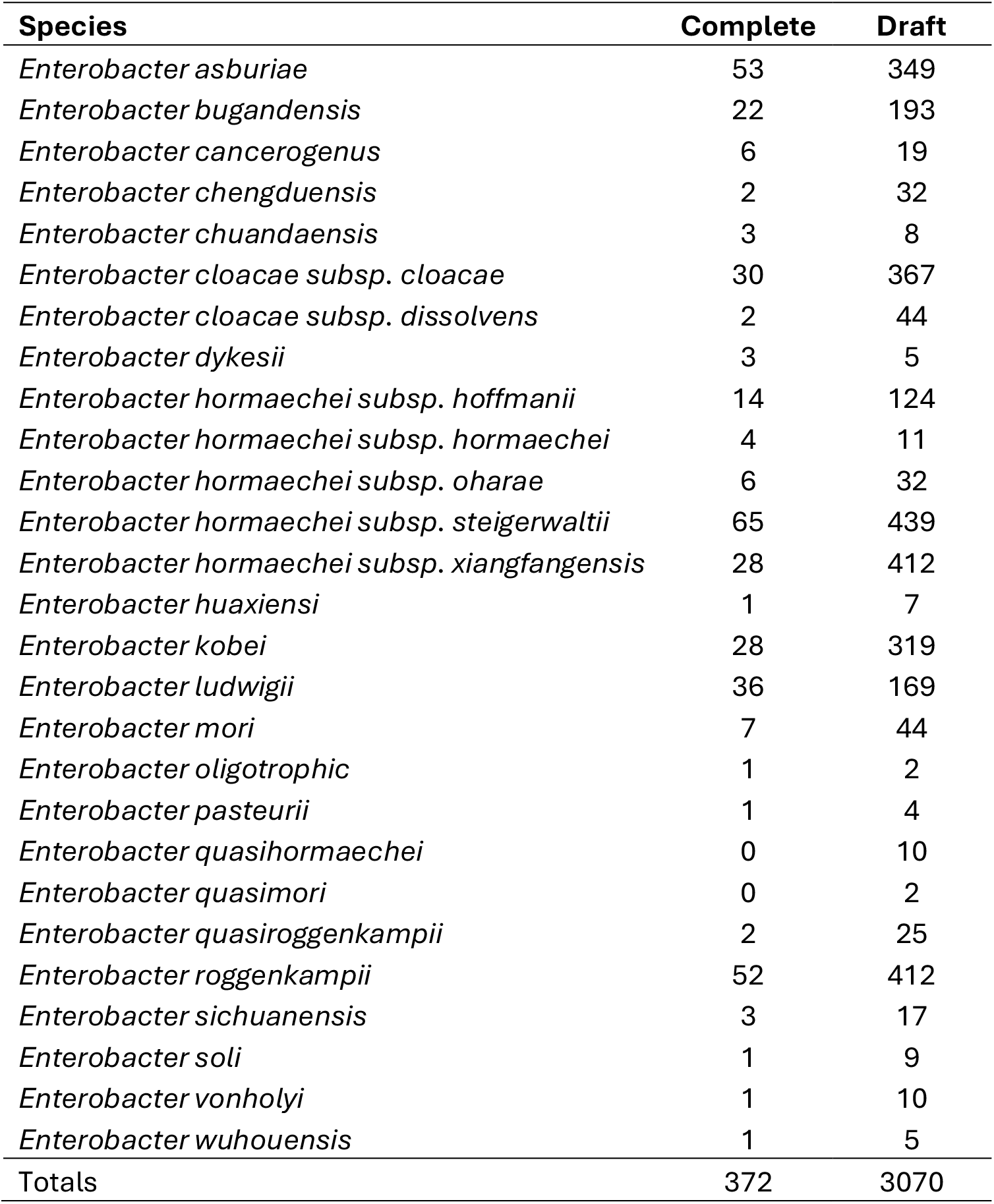
Number of draft and complete genomes from each Enterobacter cloacae complex species used to create the cgMLST schema. Genomes are listed under the species/subspecies names assigned by FastANI.

### Genomes for cgMLST schema creation

Genomes for schema creation were downloaded from the NCBI Refseq database using NCBI datasets (27). Genomes for all ECC species available on NCBI Datasets in February 2025 were downloaded, including both complete and draft assemblies but excluding atypical genomes and metagenome-assembled genomes. The only exception was genomes listed as *E. hormaechei* without a subspecies designation; because of the large number of these records, only 50 complete genomes and 500 draft genomes were downloaded. All genomes went through a quality control process to check the species ID and assembly quality. Assembly quality was checked using Quast v.5.0.2 (28) and only genomes with fewer than 150 contigs and fewer than 20 N’s per 100 kbp were retained. To verify the species ID, assemblies were compared with genomes from the type strain of each species or subspecies using FastANI version 1.33 (29). The species was assigned according to the highest ANI, using the criteria ANI > 95% for species and ANI > 98% for subspecies (12). Nineteen genomes that did not match any of the reference genomes with ANI > 95% were excluded. The 7-gene MLST sequence type from the ecloacae pubmlst scheme was assigned to each isolate using mlst v. 2.19.0 (https://github.com/tseemann/mlst). A total of 372 complete genomes and 3070 draft genomes remained after quality control and these were used to construct the schema (Table 1). Accession numbers, assembly completeness, species identified by FastANI and ST type for all samples are listed in Table S1.

### Creation of the cgMLST schema

The cgMLST schema was created using Chewbbaca v. 3.2.0 (30). A Prodigal (31) training file was created using the *E. cloacae* reference genome (isolate 1382, Refseq Accession GCF_905331265.2). This file was then used to create a whole genome MLST scheme from the 372 complete genomes using the “CreateSchema” operation in Chewbbaca (minimum Blast score ratio 0.6). “AlleleCall” was then performed to call the alleles at each locus for the complete genomes, and “ExtractCgMLST” was used to find the set of loci comprising the core genome. This initial cgMLST schema contained 2861 loci at the 95% presence threshold. CgMLST alleles were then called for the 3070 draft genomes, and the cgMLST was redetermined using the JoinProfiles and ExtractCgMLST operations.

### cgMLST clustering

Hierarchical clustering of all 3442 genomes used to create the cgMLST scheme was performed with pHierCC v. 1.24 (32), (https://github.com/zheminzhou/pHierCC) using cgMLST profiles. To determine which cluster threshold best corresponds to the division of isolates based on (a) 7-gene MLSTs and (b) species (assigned by FastANI), adjusted rand scores were calculated for every 10^th^ HierCC (HC) level using the scikit-learn python package adjusted_rand_score.

Because of the discrepancy between the currently recognised valid species names and the species proposed on the basis of WGS for *E. cloacae* and *E. hormaechei*, we chose to use the WGS-based species divisions proposed by Wu et al., 2020 and Doijad et al., 2023 (6, 10) for this analysis, as this is likely to better reflect the divisions in the cgMLST data. Therefore, isolates from *E. cloacae* subsp. *cloacae* and subsp. *dissolvens* were classed as distinct species, as were *E. hormaechei* subsp. *hormaeche*i, and *E. hormaechei* subsp. *hoffmannii*. Mean FastANI values between these pairs of taxa are close to 95% (see Table S5), in line with the accepted cutoff for species delineation of 95-96% (29). Isolates from the 3 remaining *E. hormaechei* subspecies (*oharae, steigerwaltii* and *xiangfangensis*) were grouped together as one species, as fastANI values between isolates of these taxa were above 97%.

### Evaluation of the cgMLST scheme

The schema was evaluated on two datasets known to contain epidemiologically linked isolates. The first dataset is comprised of 25 *E. cloacae* subsp. *cloacae* isolates collected in New Zealand from 2018-2024, including 19 isolates from a hospital ward outbreak collected over a 3-week period in 2024. These isolates were sent to the New Zealand Institute for Public Health and Forensic Science (PHF Science, formerly the Institute of Environmental Science and Research (ESR), Porirua, New Zealand) for whole genome sequencing as part of regular surveillance. Bacterial cultures were grown Tryptic Soy Agar (Fort Richard, Auckland, NZ) and incubated overnight at 35°C. Genomic DNA was extracted from a single colony pick using the Chemagic 360 (Perkin Elmer). Libraries were prepared using the plexWell Library Preparation kit (SeqWell) and sequenced using Illumina paired-end 150 bp sequencing on the NextSeq 2000 platform. Sequence quality checks, species identification and de novo assembly were performed using an inhouse pipeline comprising Fastp v. 0.20.1 (33), Centrifuge v.1.0.4 (34), Skesa v. 2.3.0 (35) and Quast v. 5.0.2 (28).

The second dataset comprises 27 isolates of *Enterobacter hormaechei (*subsp. *steigerwaltii, hoffmanii, and oharae)* published by Wendel et al. 2022. Isolates were from screening swabs collected from the neonatal intensive care unit (NICU) and paediatric (and neonatal) intensive care unit (PICU) in a hospital in Germany. Illumina reads were downloaded from the NCBI short read archive (project number PRJEB46479). Sequence quality checks, species identification and de novo assembly were performed using an inhouse pipeline comprising Fastp v. 0.20.1 (33), Centrifuge v.1.0.4 (34), Skesa v. 2.3.0 (35) and Quast v. 5.0.2 (28). Of the original 32 isolates in the dataset, 5 were excluded as they did not meet quality control thresholds of depth > 20 and contigs < 150.

Allele calling was performed using Chewbbaca v. 3.2.0, selecting the scheme with 99% locus-sharing. A pairwise cgMLST distance matrix was created from the allele call table produced by Chewbbaca using cgMLST-dists v.0.4.0 (https://github.com/tseemann/cgMLST-dists), and neighbour-joining and minimum-spanning trees based on the cgMLST allelic profiles were created using GrapeTree v2.1 (36). For the *E. cloacae* outbreak isolates, core genome SNP analysis was performed using Snippy v.4.6.0, using a long read assembly of the index case from this outbreak as the reference.

## Results

### Core genome MLST schema creation and evaluation across ECC species

The cgMLST schema created from 372 complete genomes and 3070 draft genomes from all *E. cloacae* complex species contains 2709 loci at the 95% loci presence threshold (i.e. loci present in 95% of genomes), and 1812 loci at 99% presence threshold and 65 loci at the 100% threshold. Having selected loci that are well represented across the whole dataset, we next assessed how completely individual genomes and specific species and subspecies were represented in the 95% vs 99% schemes to determine which had the best coverage across all ECC species. For the 95% scheme (2709 loci), 97.8% of isolates had 95% of the target loci, and 76% of isolates had 99% of the targets. However, isolates with fewer than 95% or 99% of the 2709 target loci were not evenly distributed across species: the 77 isolates with fewer than 95% of the targets came from 9 of the 27 species or subspecies, including 14% of *E. hormaechei* subsp. xiangfangensis and 21% of *E. hormaechei* subsp. *oharae*, and there were some species where 100% of isolates had fewer than 99% of the target loci (see Figure S1). For the 99% scheme with 1812 loci, 99.9% of isolates had 95% or more of the cgMLST targets, and 95.6% had 99% or more. The 3 isolates that had fewer than 95% of the gene targets were from different species: one *E. hormaechei subsp. xiangfangensis*, one *E. asburiae*, and one *E. kobei*. Because of the likelihood of species-specific missing targets with the 95% scheme, we chose to use the 99% scheme containing 1812 loci for all subsequent analyses. The coordinates of the gene targets in this scheme are given in Table S2.

We confirmed the species ID of all samples used in this study by employing the established ANI cutoff of >95% for species and >98% for subspecies (10). Where two taxa were above the threshold, the one with the highest ANI was assigned. The species or subspecies ID determined by ANI did not match the species or subspecies designated on NCBI datasets for 120 out of 3442 isolates (14 species and 106 subspecies), see Tables S3 and S4. In addition, 349 *E. cloacae* were not assigned a subspecies on NCBI datasets; 8 of these matched a different *Enterobacter* species and 37 matched *E. cloacae* subsp. *dissolvens* in the ANI analysis. A total of 535 *E. hormaechei* were not assigned a subspecies on NCBI and of these, only 9 matched *E. hormaechei* subsp. *hormaechei* with the remainder split across the other 4 subspecies. For all subsequent analysis, the taxon assigned by FastANI was used.

Analysis of cgMLST profiles show there is a high level of diversity both within and between species. There were 3144 different cgMLST profiles among 3442 genomes and a minimum-spanning tree constructed from cgMLST allele profiles shows that profiles largely cluster by species or subspecies (Figure 1). CgMLST allele distances (AD) ranged from 1402 to 1804 (mean 1782) for within-species comparisons, and 0-1804 (mean 1556) for within-subspecies comparisons. For comparisons between species, the AD ranged from 1519-1811 (mean 1801).

**Figure 1.**
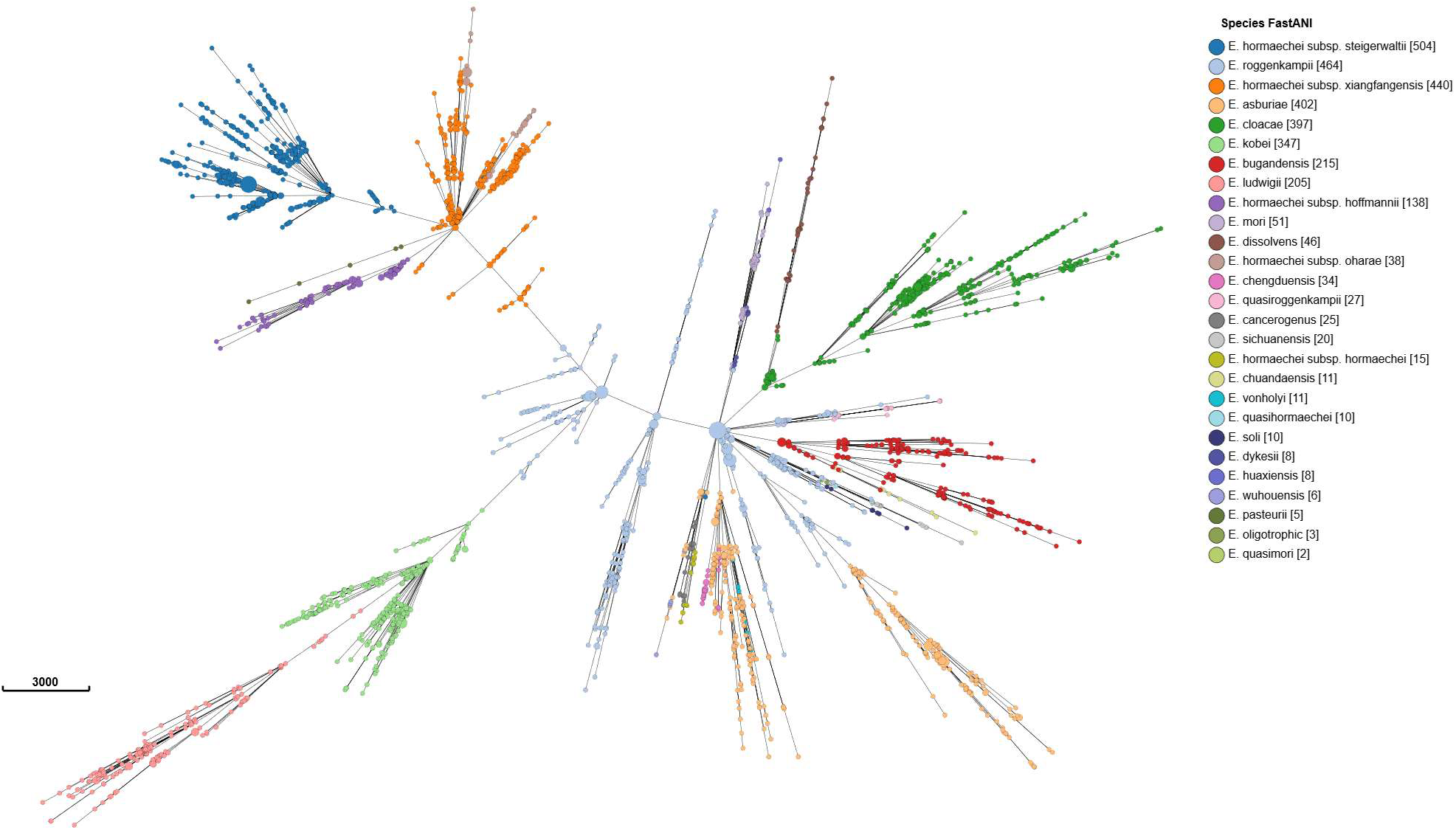
Minimum-spanning tree of cgmlst profiles for all 3442 isolates used to create and refine the cgMLST schema. Nodes are coloured according to the species or subspecies assigned by FastANI comparisons to type species.

Adjusted rand index (ARI) was calculated between MLST STs, species ID and cgMLST clusters at every 10^th^ hierCC level to determine which HC level provides clustering most concordant with division of isolates by MLST or species assignment. There was almost perfect concordance (ARI=0.948) between MLST and cgMLST at HC350, and strong concordance (ARI=0.883) between species divisions and cgMLST at HC1730 (Figure S2). At HC1730, samples fell into 64 clusters, of which 16 only contained a single isolate (referred to as singletons from here on). Many species were represented by a single cgMLST cluster which was unique to that species (see Table 2 and Figure S3). Two clusters contained isolates from more than one taxon: one that contained isolates from *E. hormaechei* subsp. *xiangfangensis*, subsp. *oharae*, subsp. *xiangfangensis*, which was expected as these were considered one species in the ARI analysis; and another contained isolates from *E. asburiae* and *E. roggenkampii*. Further analysis of *E. asburiae* and *E. roggenkampii* hierCC levels showed that species-specific clusters for these two species were obtained at hierCC level 1669.

**Table 2.** Comparison between number of MLST STs, cgmlst STs (cgSTs), and cgmlst clusters (containing more than 1 isolate) at HC350 and HC1730 for all Enterobacter cloacae complex species.

| Species | #<br>genomes | # ST<br>types | #cgST<br>types | #cgmlst<br>clusters<br>HC350 | # cgmlst<br>clusters<br>HC1730 | #cgmlst<br>singletons<br>HC1730 |
| --- | --- | --- | --- | --- | --- | --- |
| *E. asburiae* | 402 | 104 | 364 | 137 | 10 | 6 |
| *E. bugandensis* | 215 | 89 | 202 | 109 | 2 | 0 |
| *E. cancerogenus* | 25 | 6 | 25 | 16 | 1 | 1 |
| *E. chengduensis* | 34 | 6 | 32 | 5 | 2 | 1 |
| *E. chuandaensis* | 11 | 4 | 11 | 7 | 1 | 0 |
| *E. cloacae* subsp. *cloacae* | 397 | 91 | 366 | 111 | 1 | 0 |
| *E. cloacae* subsp. *dissolvens* | 46 | 30 | 46 | 39 | 1 | 0 |
| *E. dykesii* | 8 | 3 | 6 | 5 | 1 | 1 |
| *E. hormaechei* subsp.<br>*hoffmannii* | 138 | 30 | 130 | 27 | 1 | 0 |
| *E. hormaechei* subsp.<br>*hormaechei* | 15 | 4 | 15 | 5 | 1 | 0 |
| *E. hormaechei* subsp. *oharae* | 38 | 8 | 28 | 6 | 1 | 0 |
| *E. hormaechei* subsp.<br>*steigerwaltii* | 504 | 97 | 458 | 100 | 2 | 1 |
| *E. hormaechei* subsp.<br>*xiangfangensis* | 440 | 76 | 411 | 71 | 1 | 0 |
| *E. huaxiensis* | 8 | 0 | 7 | 5 | 1 | 0 |
| *E. kobei* | 347 | 77 | 330 | 78 | 1 | 0 |
| *E. ludwigii* | 205 | 66 | 192 | 101 | 1 | 0 |
| *E. mori* | 51 | 9 | 50 | 41 | 4 | 3 |
| *E. oligotrophic* | 3 | 0 | 3 | 2 | 1 | 0 |
| *E. pasteurii* | 5 | 2 | 5 | 4 | 1 | 0 |
| *E. quasihormaechei* | 10 | 1 | 10 | 7 | 3 | 1 |
| *E. quasimori* | 2 | 0 | 2 | 2 | 1 | 0 |
| *E. quasiroggenkampii* | 27 | 12 | 27 | 24 | 1 | 0 |
| *E. roggenkampii* | 464 | 135 | 377 | 149 | 2 | 1 |
| *E. sichuanensis* | 20 | 12 | 20 | 17 | 2 | 0 |
| *E. soli* | 10 | 2 | 10 | 7 | 2 | 0 |
| *E. vonholyi* | 11 | 10 | 11 | 10 | 2 | 0 |
| *E. wuhouensis* | 6 | 0 | 6 | 6 | 1 | 1 |
| **Totals** | **3442** | **874** | **3144** | **1091** | **48** | **16** |

### Performance of the cgMLST schema for identifying outbreaks

To investigate the suitability of the cgMLST scheme for identifying isolates in an outbreak, isolates from two datasets were typed. The first dataset contains 25 *E. cloacae* subsp. *cloacae* isolates collected in New Zealand from 2018-2024, including 19 isolates from a hospital-associated outbreak with known epidemiological linkages (Figure 2). The outbreak isolates form a single cluster with cgMLST allele distances ranging from zero to three (mean pairwise cgmlst AD= 0.5, mode = 0). Slightly higher distances were observed with core genome SNP analysis, with 0-4 SNP differences between isolates in the cluster (mean pairwise SNPs = 1.7, mode 2). The other 6 isolates were divergent from each other and the outbreak isolates with allele difference values ranging from 669-1724 (mean 1707.2).

**Figure 2.**
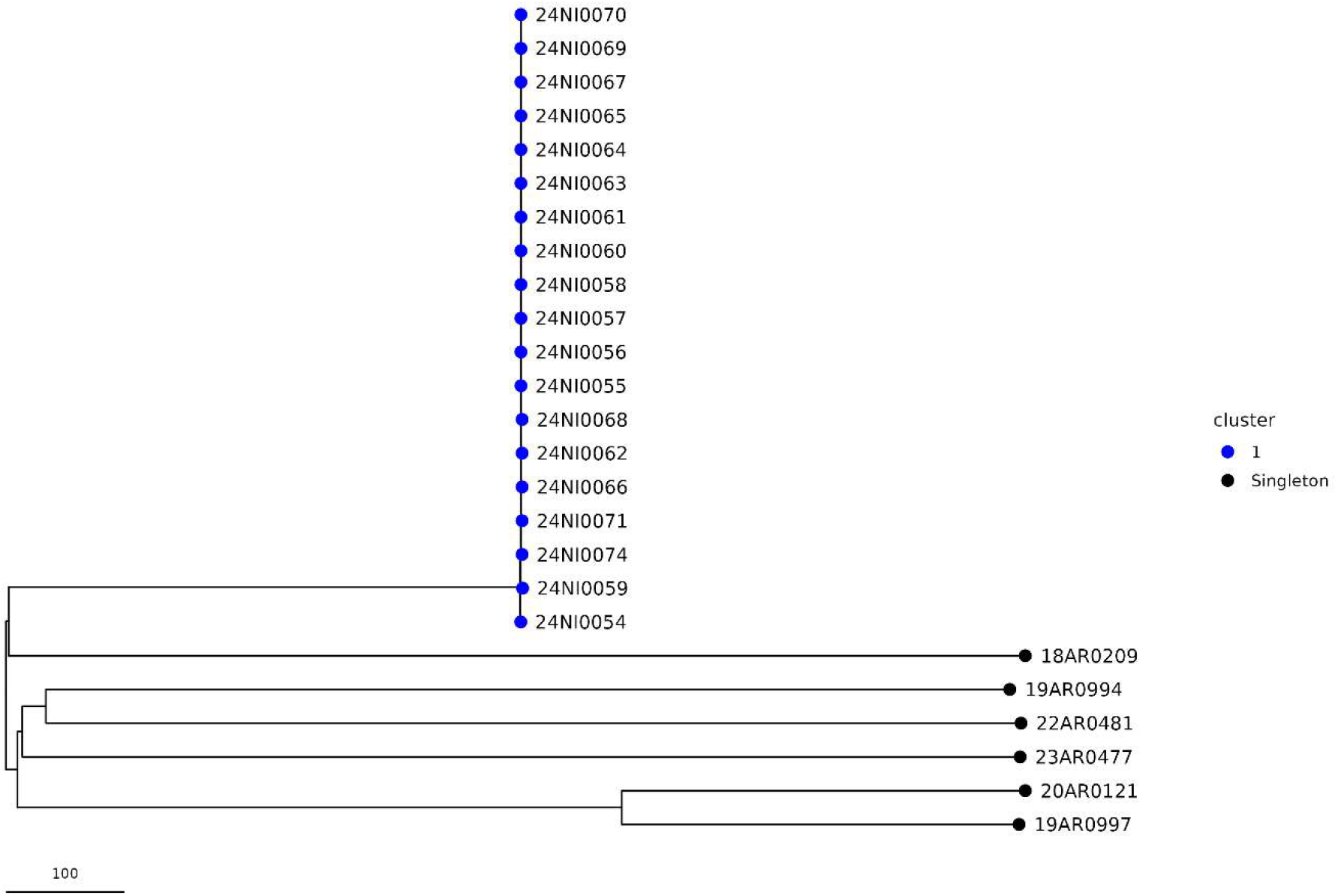
Neighbour-joining tree of cgmlst profiles for E. cloacae subsp. cloacae isolates collected in New Zealand from 2018-2024. Nodes are coloured by cgMLST cluster with a cluster threshold of 5 alleles.

The second dataset comprised 27 isolates from a neonatal intensive care unit in Germany, collected over a 6 month period in 2017-2018 and published by Wendel et al in 2022 (1). The cgMLST typing grouped these isolates into 4 clusters, plus 3 single isolates (Figure 3). Within each cluster, isolates were separated by 0-3 allele differences (mean within-cluster allele difference 1.3). For singleton and between-cluster comparisons allele difference values ranged from 1625 to 1800 (mean 1770.1). The four clusters identified by cgMLST were identical to those identified by Wendel et al. using core genome SNP analysis, PFGE and RAPD typing. Isolates within each cluster were confirmed to have epidemiological links and were the same ST type. Wendel et al. measured 0-1 SNP differences within each cluster, and at least 180 SNPs between clusters.

**Figure 3.**
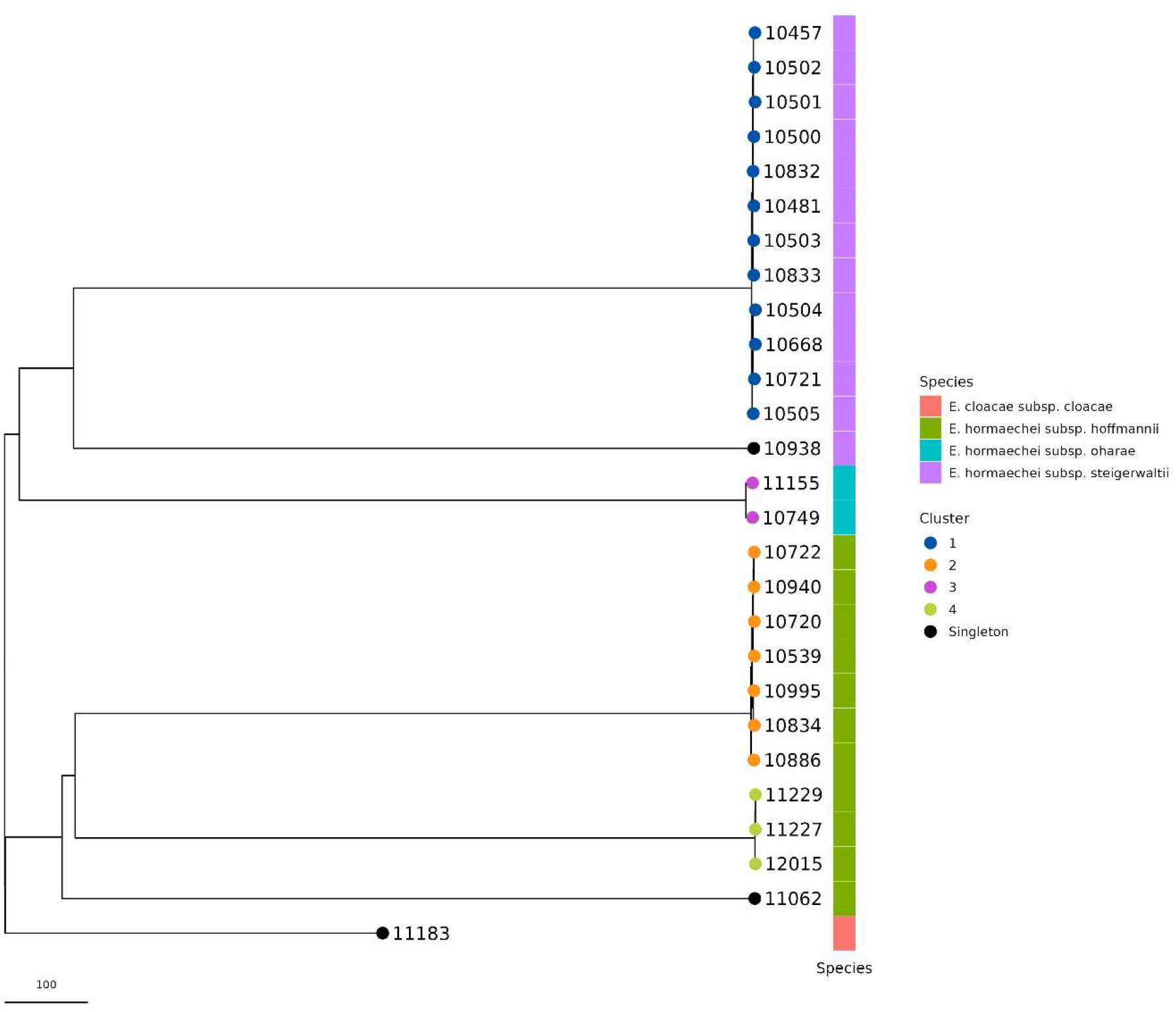
Neighbour-joining tree of cgmlst profiles for ECC isolates from a neonatal unit in Germany. Nodes are coloured by cgMLST cluster with a cluster threshold of 5 alleles, and species is shown in the coloured bar.

## Discussion

The development of a cgMLST scheme for the *E. cloacae* species complex represents a critical step toward standardized genomic epidemiology of this clinically important group. Because accurate species identification can be challenging for this group, there is a need for a surveillance tool that does not require accurate identification of the species by laboratory methods first, and that can accommodate any future taxonomic revisions. When checking the species assigned to each genome in NCBI datasets against ANI comparisons to type strains we found that 3.5% of samples were assigned to a different species or subspecies by ANI compared with NCBI, and 77% and 47% of *E. cloacae* and *E. hormaechei* respectively did not have a subspecies assigned on NCBI. The latter is particularly problematic due to the debate around whether *E. cloacae subsp. dissolvens* and *E. hormaechei* subspp. *xianfangensis* and *hoffmannii* should be elevated to full species (6, 10). Although ANI should not be the sole determinant of species, these discrepancies underscore the need for a typing scheme that works across the species complex. By using a large collection of 3,442 genomes from 27 species or subspecies we were able to define a cgMLST scheme that works equally well across all species within the complex yet retains sufficient resolution to be able to trace outbreaks. The cgMLST schema provides a genomic framework that is robust to future taxonomic revisions without loss of comparability, as loci are defined at the core-genome level rather than being tied to a particular taxonomic assignment.

The scheme comprises 1812 target loci at the 99% loci presence threshold and 2706 loci at the 95% threshold. Testing with almost all genomes available on NCBI showed that the scheme with 1812 loci was more broadly representative for all ECC species, as 99.9% of isolates had more than 95% of the target loci, and only single isolates from 3 different species had fewer than 95% of loci. In contrast, the scheme with 2706 loci had more uneven coverage across species. Although 97.8% of isolates overall had more than 95% of the target loci, nine taxa contributed the majority of isolates with fewer than 95% of targets, including the clinically important *E. hormaechei* subspecies *xiangfangensis* (14 % of its isolates) and *E. hormaechei* subspecies *oharae* (21 % of its isolates). Thus, the 99% scheme with 1812 targets better represents the core genome across all ECC species, so we proceeded with this schema for subsequent analyses.

Although this scheme was developed to support cluster identification and public health surveillance, we investigated how well cgMLST-based clustering recapitulates known species and subspecies divisions within the complex. In addition to comparing to current taxonomy of the group, this analysis provides an indirect measure of the schemes ability to capture deeper population structure. Across the genomes used in this study, cgMLST profiles largely cluster by species or subspecies and reflect the expected phylogenetic structure of the species complex, despite the overall high levels of diversity in cgMLST profiles both within and between species. To investigate whether cgMLST could be used to differentiate between species within the complex, we used hierarchical clustering to determine which cluster threshold most closely matched the division of isolates into species based on FastANI thresholds. Hierarchical clustering allows analysis of isolates at a wide range of cluster thresholds in a single step, where higher levels may be suitable for defining lineages at species level within the species complex, and lower levels provide greater resolution for determining relationships within putative outbreak clusters ((32),(17),(37)). Using Adjusted Rand Index, we determined that hierCC level 1730 had highest concordance with species divisions. Of the 27 species or subspecies included in this study, 22 had species-specific cgMLST clusters at this threshold and for 17 taxa, all samples within the taxon were contained within a single, specific, cgMLST cluster. We observed one instance of isolates from 2 different species falling into a single cluster at this threshold, but these species could be separated at a lower hierCC threshold (HC1669). It should be noted that ability to draw firm conclusions is limited by uneven representation of species within our dataset, with some species represented by fewer than 10 genomes, and sampling that may not have been random. Nevertheless, these results suggest that systematic allele numbering and cgmlst profile codes could, in the future, be used for species identification or the creation of stable LIN codes to support classification with the ECC.

Analysis of two datasets containing samples with known epidemiological links and core genome SNP data for comparison demonstrated that, although diversity is high in unrelated strains, the scheme is suitable for identifying closely related isolates. In the New Zealand *E. cloacae* subsp. *cloacae* outbreak, outbreak isolates differed by 0–3 cgMLST alleles (mean 0.5) and 0–4 SNPs (mean 1.7), indicating tight concordance between allele- and SNP-based inferences. On the German NICU dataset (*E. hormaechei* subspp. *steigerwaltii, hoffmannii, oharae*), our cgMLST analysis defined four clusters with 0–3 allele differences within clusters and ≥1,600 alleles between clusters, mirroring the clusters identified by WGS-SNP phylogenies in the original study. These results are in line with studies on other species showing that cgMLST has similar discriminatory power to core genome SNP analyses for detecting outbreaks (16, 24, 38). As all outbreak-linked isolates differed by fewer than 3 alleles in both studies we suggest this is an appropriate threshold for interfering recent transmission. However, a threshold of 5 alleles would allow higher sensitivity, for example when tracking outbreaks over a longer time period, with the usual caveat that thresholds should be interpreted alongside epidemiology and sampling context. Putative clusters identified using cgMLST can be followed up with SNP typing to provide additional resolution.

In summary, our ECC cgMLST scheme bridges the gap between the high phylogenetic resolution of SNP typing and the portability and scalability required by public health practice. It performs consistently across a taxonomically challenging species complex, aligns closely with core genome SNP typing for outbreak-scale relationships, and provides a practical framework for standardized ECC genomic epidemiology.

