## Supplementary figures for "Development and evaluation of a core genome multi-locus sequence typing scheme for the *Enterobacter cloacae* complex"

Supplementary Figure 1. Percent of cgMLST gene targets across all species for 95% (2709 targets) vs 99% (1812 targets) schema.

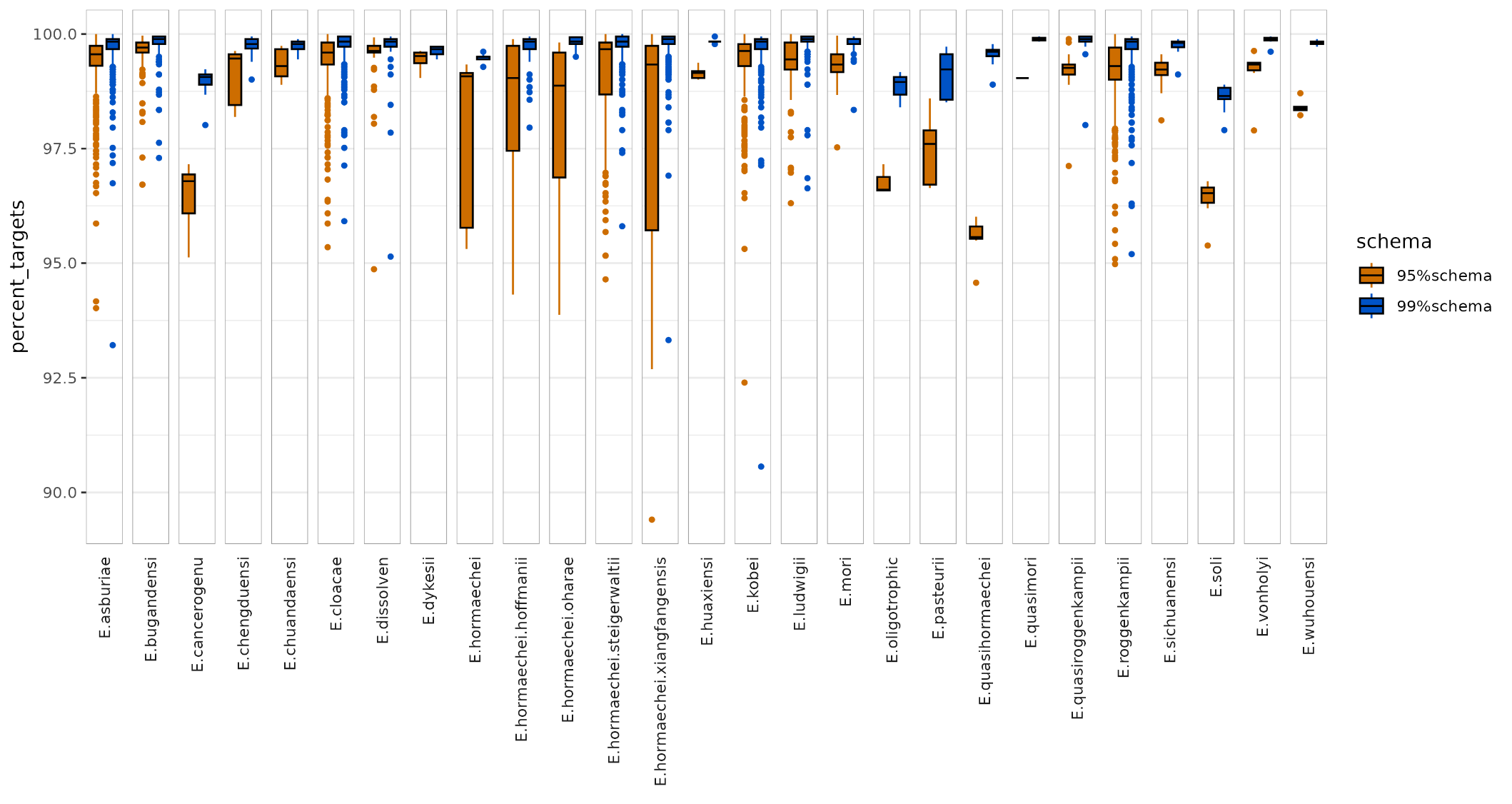

Supplementary Figure 2. Adjusted Rand Index (ARI) comparison between clusters in each HierCC level with (A) MLST STs and (B) species. The blue line indicates ARI between clusters at every 10^th^ hierCC level.

A.

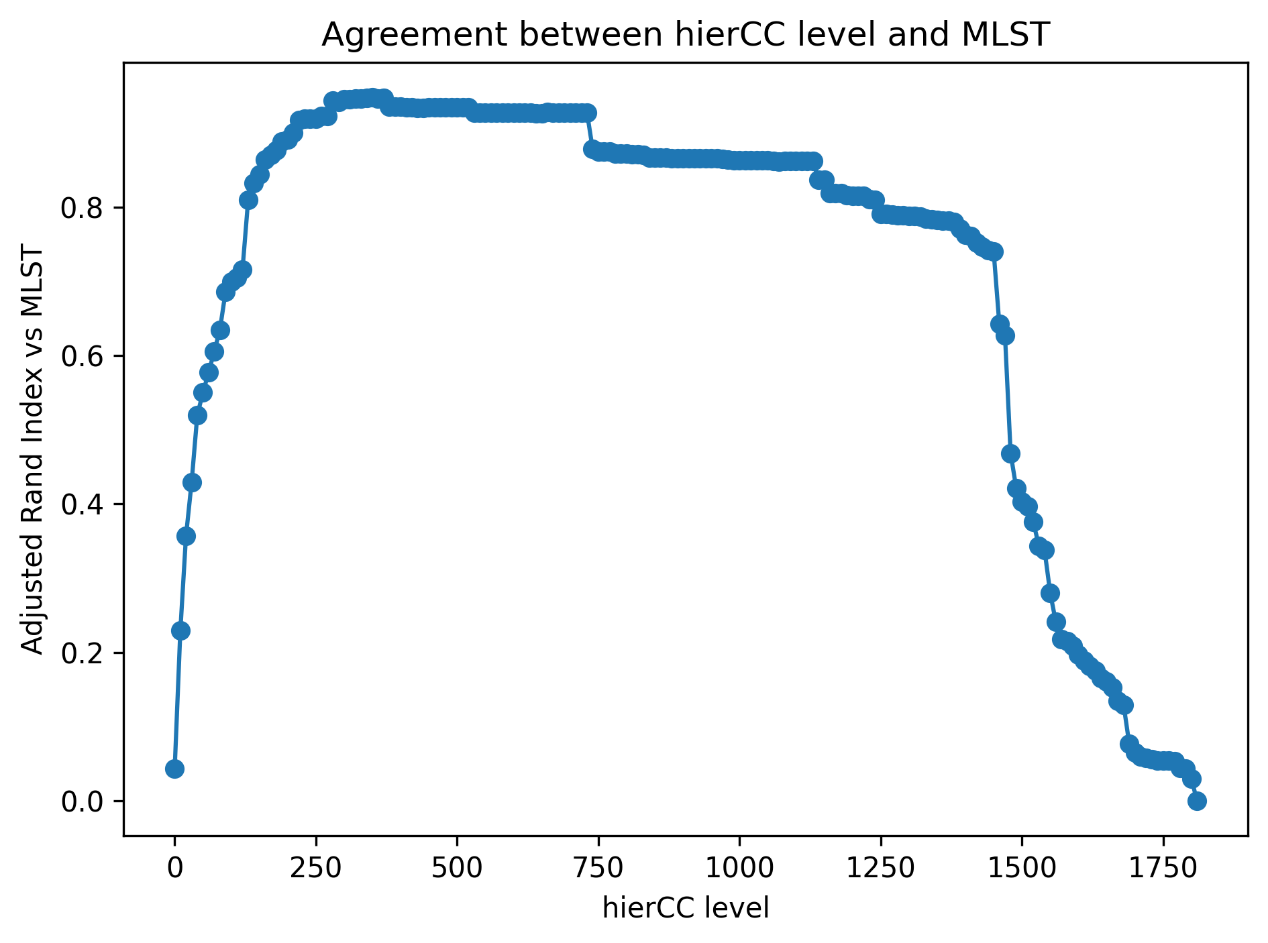

B.

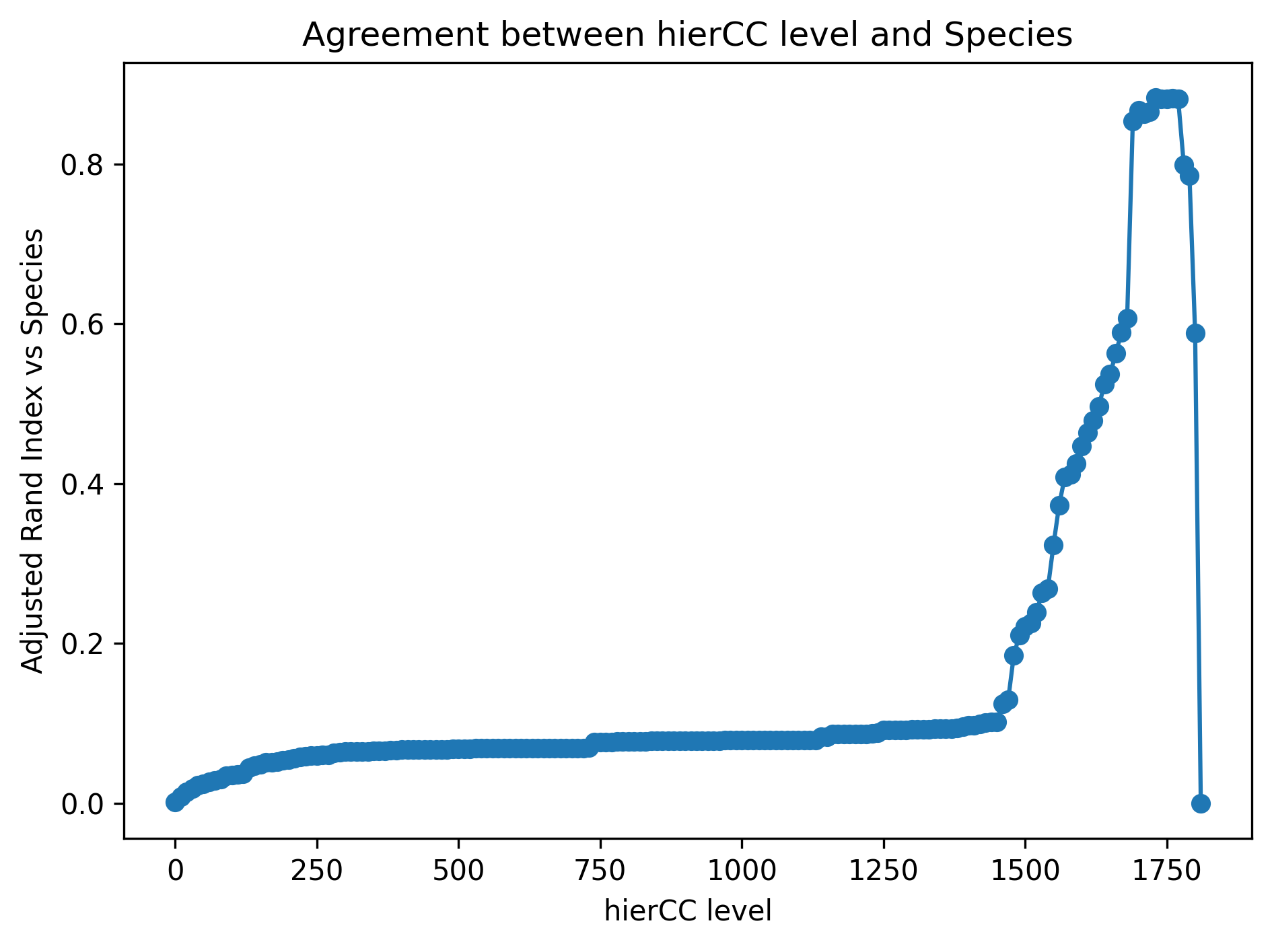

Figure S3. Neighbour-joining tree of cgMLST profiles for all 3442 isolates used to create the cgMLST scheme, showing the concordance between species (1^st^ column) and cgMLST cluster at hierCC level 1730 (2^nd^ column).

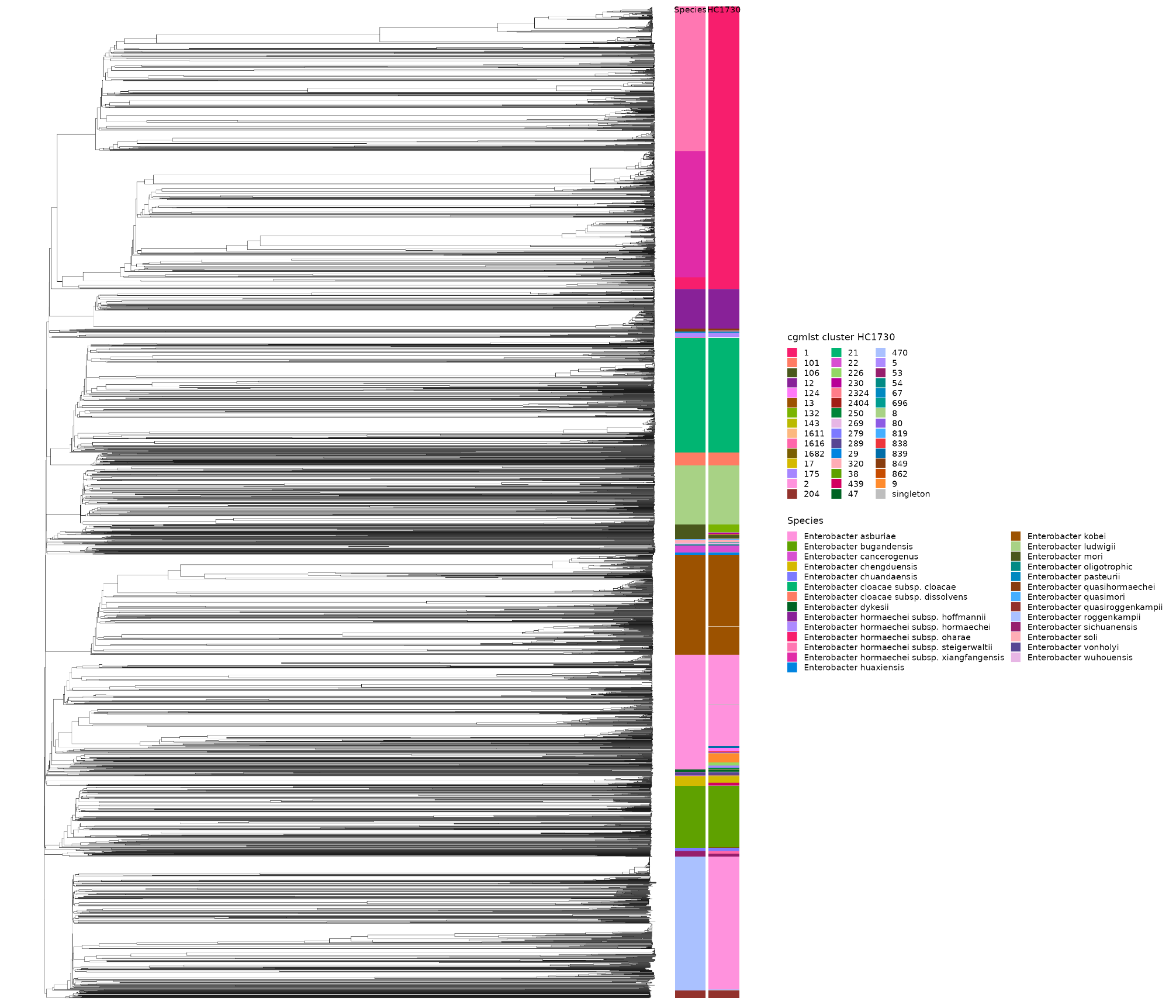

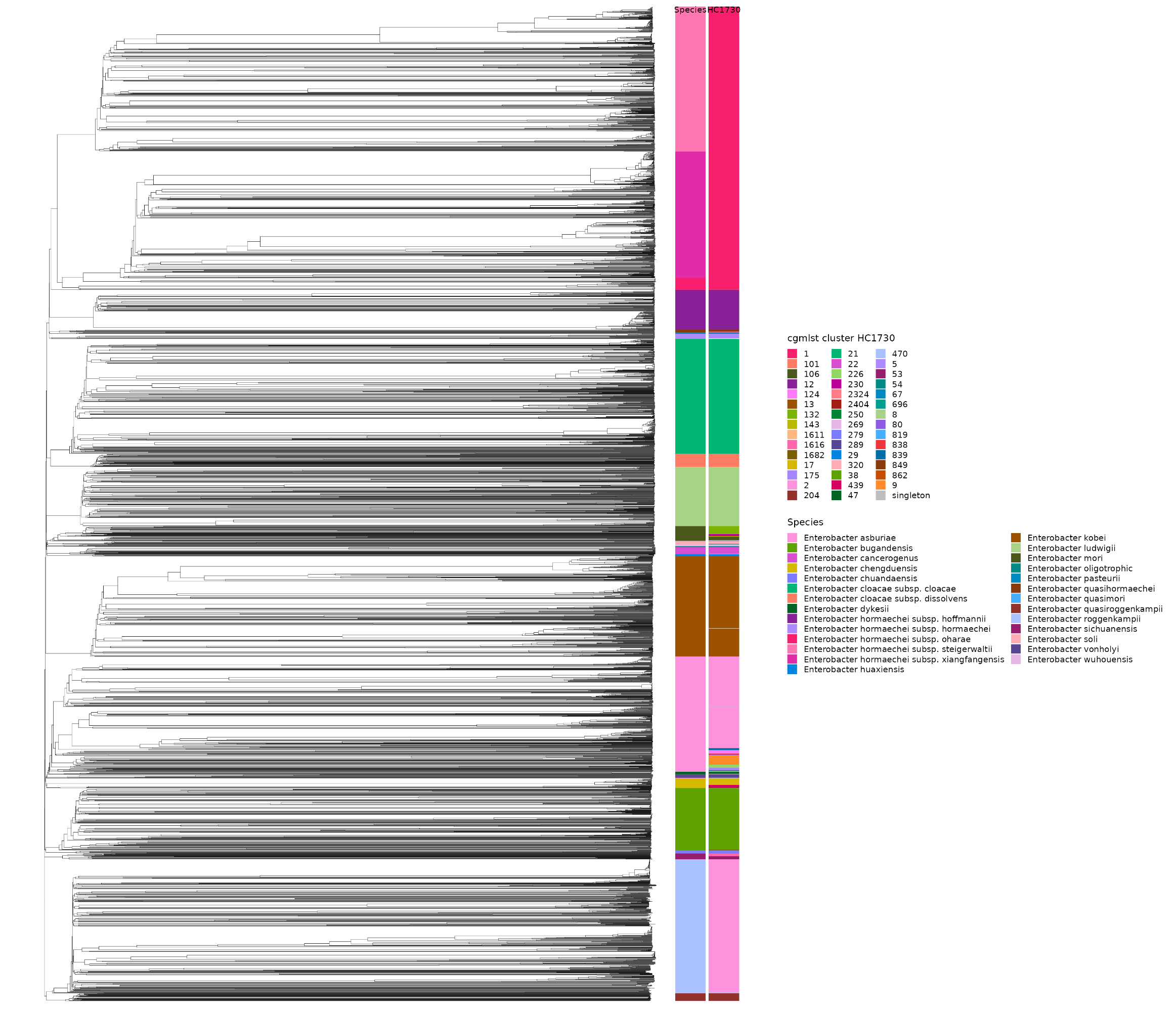

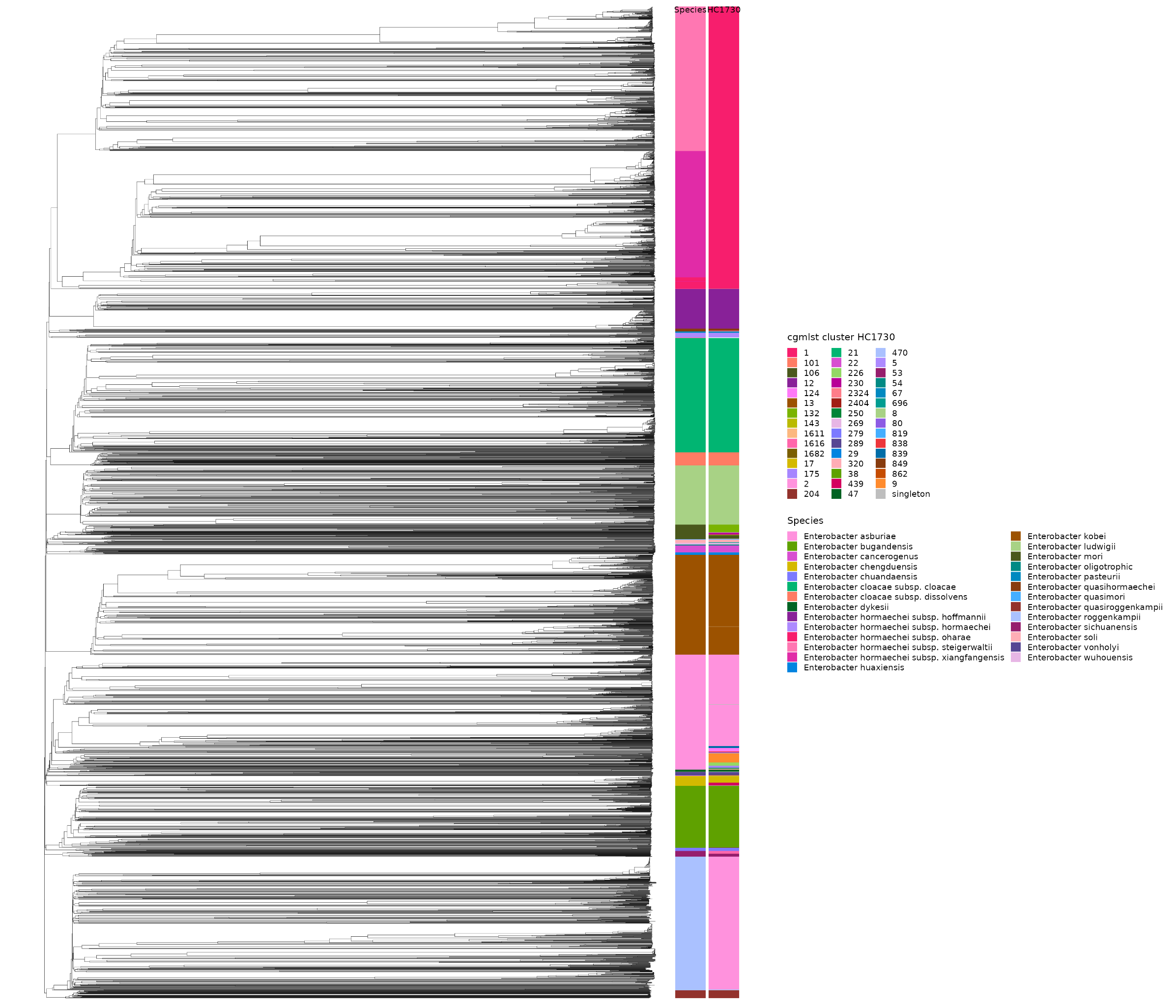
